# Inositol synthesis gene is required for circadian rhythm of *Drosophila melanogaster* mating behavior

**DOI:** 10.1101/2020.02.19.955583

**Authors:** Kazuki Sakata, Haruhisa Kawasaki, Norio Ishida

**Author notes:** Corresponding author: Norio Ishida Institute for Chronobiology, Foundation for Advancement of International Science, 3-24-16 Kasuga, Tsukuba, Ibaraki, 305-0821, Japan., (Norio Ishida), (Norio Ishida).

## Abstract

Accumulating evidence indicates that the molecular circadian clock underlies the mating behavior of *Drosophila melanogaster.* However, information about which gene affects circadian mating behavior is poorly understood in animals. The present study found that feeding *Myo*-inositol enhanced the close-proximity (CP) rhythm of *D. melanogaster* mating behavior and lengthened the period of the CP rhythm. Then, to understand a role for inositol synthesis to fly mating behavior, we established the *Inos* (*Myo*-inositol 1-phosphate synthase) *gene* knock down fly strains with RNAi. Interestingly, the CP behavior of this three-different driver knock down strains was arrhythmic, but the locomotor rhythm was rhythmic. The data of three-different *Inos* knock down strains suggests that *Inos* gene expression of upper LN_d_, l-LN_V_, 5^th^s-LN_v_ in brain is necessary for proper CP rhythm generation in *D. melanogaster*. The data indicated that the *Inos* gene is involved in the role for the circadian rhythm of *D. melanogaster* mating behavior.

## 1. Introduction

The physiology and behavior of many organisms allows adaptation to daily and seasonal environmental changes via circadian clocks that comprise an endogenous self-sustained timekeeping system (Dunlap, 1999; Ishida, Kaneko, & Allada, 1999). Furthermore, the molecular mechanisms of circadian clock genes that comprise transcriptional–translational feedback loops are conserved from flies to humans (Kako & Ishida, 1998). A core oscillator mechanism of circadian rhythms and feedback loops involving several clock genes such as *period* gene (Konopka & Benzer, 1971), control the locomotor activity and eclosion of the fruit fly, *Drosophila melanogaster* (Dunlap, 1999). The relationships between behavioral rhythms and circadian clock genes have been studied in mutants of the fly with defective clock gene. The circadian rhythm of mating behavior is controlled by the clock genes, *period* and *timeless* in *D. melanogaster* (Sakai & Ishida, 2001). Accumulating evidence indicates that the circadian clock underlies the reproductive behavior of *D. melanogaster* (Beaver & Giebultowicz, 2004; Kadener et al., 2006). Heterosexual fly couples exhibit significantly di□erent circadian activity from individual flies, having a brief rest phase around dusk followed by activity throughout the night and whole day. This activity between heterosexual fly couples is referred to as the close-proximity (CP) rhythm (Fujii, Krishnan, Hardin, & Amrein, 2007; Hamasaka, Suzuki, Hanai, & Ishida, 2010). Analyses of CP rhythms have shown that circadian clocks regulate male courtship behavior in a circadian manner and that a core component of the circadian clock gene, *per*, is needed to generate CP rhythms. We and others reported the brain clock neurons that are required for CP rhythms (Fujii et al., 2007; Hamasaka et al., 2010).

*Myo*-inositol, formerly referred to as vitamin B7, is a cyclitol naturally occurring in nature and foodstuffs. *Myo*-inositol exists under nine distinct stereoisomers (named *allo*-, D-*chiro*-, L-*chiro*-, *cis*-, *epi*-, *muco*-, *myo*-, *neo*- and *scyllo*-inositol) through 4 pimerization in hydroxyl group configuration. *Myo*-inositol is the important precusor for a key second messenger, *Myo*-inositol 1,4,5-triphosphate (IP3) in mammals (Sarkar & Rubinsztein, 2006). *Myo*-inositol is synthesized naturally from glucose in animals. In mice, *Myo*-iositol have shown insulin mimetic properties such as lowering postprandial blood glucose level and stimulating glucose uptake (Croze, Geloen, & Soulage, 2015). In *D. melanogaster*, our previous paper found that dietary *Myo*-inositols affect the mating circadian rhythm of CP behavior that reflects male courtship motivation in flies (Sakata et al., 2015). A recent study reported that *Myo*-inositol synthesis is involved in maintaining the period of circadian behavior in mice (Ohnishi et al., 2014). *Myo*-inositol may be beneficial for depressed patients (Mukai, Kishi, Matsuda, & Iwata, 2014; Zhao et al., 2015) and important for fungal sexual differentiation, reproduction and virulence (Niederberger et al., 1998; Voicu, Poitelea, Schweingruber, & Rusu, 2002). *Myo*-inositol is also used to increase clinical pregnancy rates in infertile men and women (Korosi, Barta, Rokob, & Torok, 2017; Regidor & Schindler, 2016). However, precisely how *Myo*-inositol is involved in the molecular mechanisms of diverse mating phenomena from fungi to humans is unknown. We tried to determine the molecular mecanism of *Myo*-inositol in the circadian mating rhythms of *Drosophila melanogaster*.

The African ice plant, Mesembryanthemum crystallinum, is abundant in *Myo*-inositols that improves hyperglycemia and memory impairments is diabetes model rat (Lee, Lee, & Wu, 2014). We previously found that powdered ice plant gradually increases the CP behavior of *D. melanogaster* under low-nutrient conditions (Sakata et al., 2015).

The present study evaluated the effects of high concentration of dietary *Myo*-inositol on the circadian rhythms of CP behavior in *D. melanogaster*. We also found the effects of internal *Myo*-inositol synthesis on CP rhythm by using the *Inos* RNAi knock down strain with three different drivers. We uncovered the importance of the *Inos* gene and its brain expression sites for the circadian mating rhythms of *D. melanogaster*.

## 2. Results

### 2.1. Dietary *Myo*-inositol lengthens the period of *Drosophila* CP and locomotor rhythms

We previously determined that the CP rhythm of *D. melanogaster* mating behavior requires the circadian clock gene, *per* (Sakata et al., 2015). Here, we aimed to determine the sex for which the CP rhythm is important. Figure 1a shows that the CP rhythm is lost if the male is a *per*^*0*^ mutant. This finding suggests that the CP rhythm in this system reflects male mating motivation behavior.

**Fig. 1.**
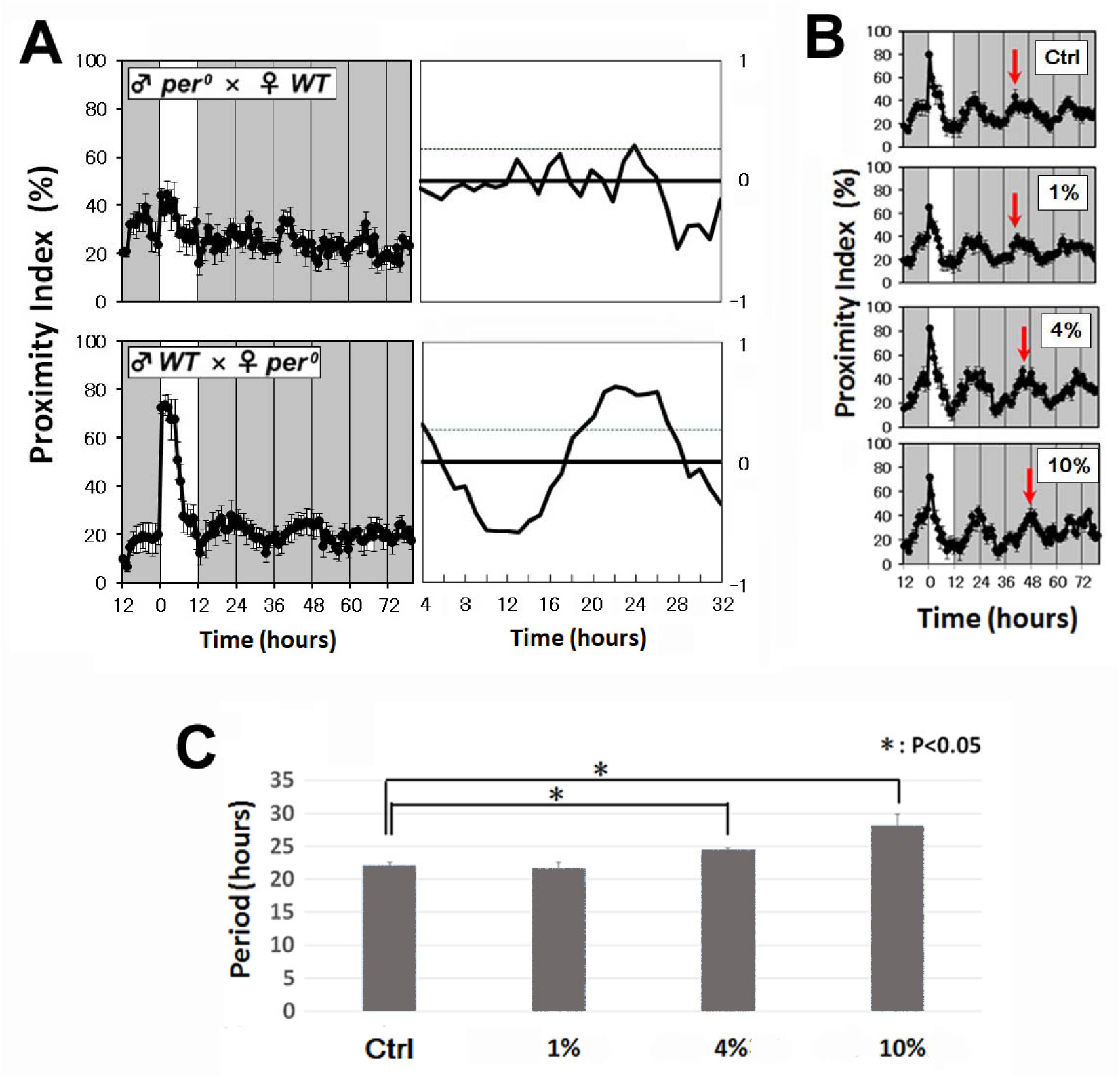
Dietary *Myo*-inositol affects CP rhythm in *Drosophila*. (A) Male plays most important role for CP rhythm in *Drosophila*. CP rhythm is abolished when male fly *per*^*0*^ mutant crossed with wild-type female. But vice-versa couple is still rhythmic (n=10 each). White area on the graph indicates day; black and gray bars indicate subjective night and subjective day, respectively. All CP rhythms were statistically tested by autocorrelation analysis (right panels of each), resulting in significant circadian rhythmicity (95% significant level indicated by dotted line). (B) The period of CP rhythm (WT × WT; n=5 each) is lengthened by *Myo*-inositol with dose-dependent manner in LNF. Arrows indicate peaks of each cycle. White area on the graph indicates day; black and gray bars indicate subjective night and subjective day, respectively. (C) The periods of CP rhythm were quantified with different *Myo*-inositol concentration.

In our previous paper, we showed that low concentration (0.001, 0.01, 0.1%)*Myo*-inositol slightly shortened the period of the CP rhythm (Sakata et al., 2015). In this paper, we tried to test high concentration effect of *Myo*-inositol to CP rhythm. Interestingly, the period of CP rhythm was longer in flies fed with a low-nutrient (LNF) diet containing 4% and 10% *Myo*-inositol than in controls (Fig. 1C). The period of the locomotor rhythm of wild-type *D. melanogaster* was also lengthened when fed with 2%, 5% and 10% dietary *Myo*-inositol (Fig. S1). These data indicated that exogenous dietary *Myo*-inositol affects the period of locomotor behavior as well as CP rhythms in *D. melanogaster*.

### 2.2. Role of the *Inos* (*Myo*-inositol 1-phosphate synthase) gene for locomotor rhythm generation

Fig. 2A shows the pathway of *Myo*-inositol synthesis in *D. melanogaster*. The *Inos* (*Myo*-inositol 1-phosphate synthase) gene is important for this synthesis step. To determine the internal synthesis effect of *Myo*-inositol on rhythmic behavior in flies, we knocked down *Inos* gene expression via RNAi with different drivers. The drivers of the three RNAi lines were *armadillo*-GAL4, *period-*Gal4 and *neuropeptide F*-Gal4. We evaluated the effects of these knock down strains on locomotor behavior (Fig. 2B) and found that the period of locomotor behavior in the three RNAi lines did not significantly differ from parental control strains (driver lines, pomotor-Gal4 lines). The data suggests that the *Inos* gene expression by these three different drivers is not required for the rhythmic locomotor behaviors.

**Fig. 2.**
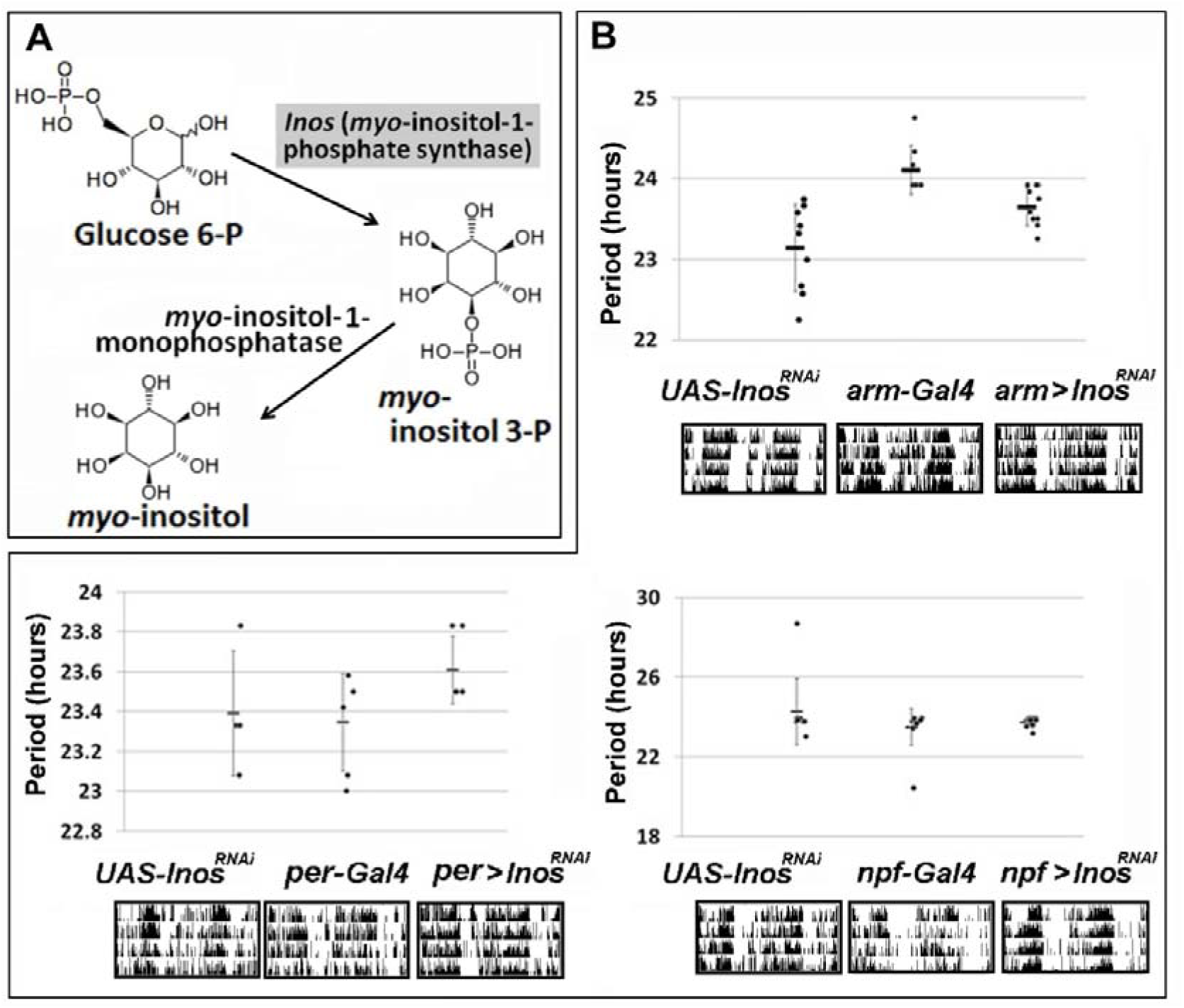
*Myo*-inositol synthesis is not required for locomotor rhythms in *Drosophila*. (A) Glucose 6-P is converted to *Myo*-inositol 3-P by *Inos* (*Myo*-inositol-1- phosphate synthase) (Park & Kim, 2004). (B) Locomotor rhythms of *UAS-Inos*^*RNAi*^ drived by the three different driver strains were compared with the parental strains (control). Data are shown as means ± SE. Detected locomotor rhythms of three different RNAi strains did not show any significant difference with two parental control lines. Actograms of each strain are shown at bottom.

### 2.3. Role of the *Inos* gene expressions for CP rhythm generation

We determined the CP rhythms of three different driver RNAi lines. Interestingly, the CP rhythm of three different driver RNAi lines were not detectable compared with both of parental control strains (driver lines, promotor-Gal4 lines) (Fig. 3C, I, O). Furthermore, autocorrelation analysis showed that the maximum values were about 24 hours in parental control strains (driver lines, pomotor-Gal4 lines). In contrast, 24 hours peaks did not detected in three different driver RNAi lines (Fig. 3F, L, R). These findings imply that internally synthesized *Myo*-inositol is necessary to generate CP rhythms in the RNAi expressed requires of *D. melanogaster*.

**Fig. 3.**
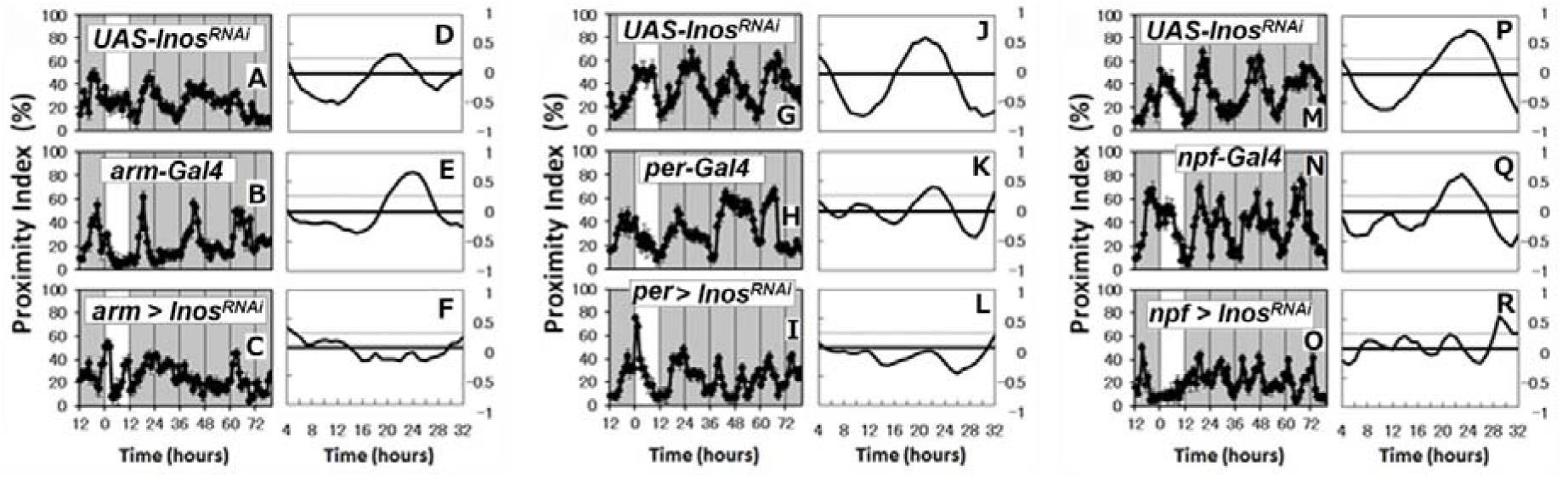
The loss of CP rhythms in three different *Inos* gene RNAi mutant strains. Although proximity index shows obvious circadian rhythms in control strains (driver lines: panels A, G, M; pomotor-Gal4 lines: panels B, H, N), these rhythms were abolished in *Inos* knock down strains: panels C, I, O (n=5 each). White area on the graph indicates day; black and gray bars indicate subjective night and subjective day, respectively. All CP rhythms were statistically tested by autocorrelation analysis (panels D-F, J-L, P-R). Autocorrelation analysis detected no circadian rhythmicity in the three *inos* knock down strains (panels F, L, R).

### 2.4. Common brain region that is important for the *Inos* gene expression in *Drosophila* CP rhythm

We attempted to identify the common brain region that is important for generating CP rhythms. We summarized the common expression region using three different promotors (Fig. 4). The fact that the CP rhythm was lost in all three RNAi lines (Fig. 3) suggests that the common brain region for the *Inos* gene RNAi expression is upper LN_d_, l-LN_V_ and 5^th^ s-LNv. The *Inos* gene expression in these three regions might be important for *D. melanogaster* CP rhythm generation.

**Fig. 4.**
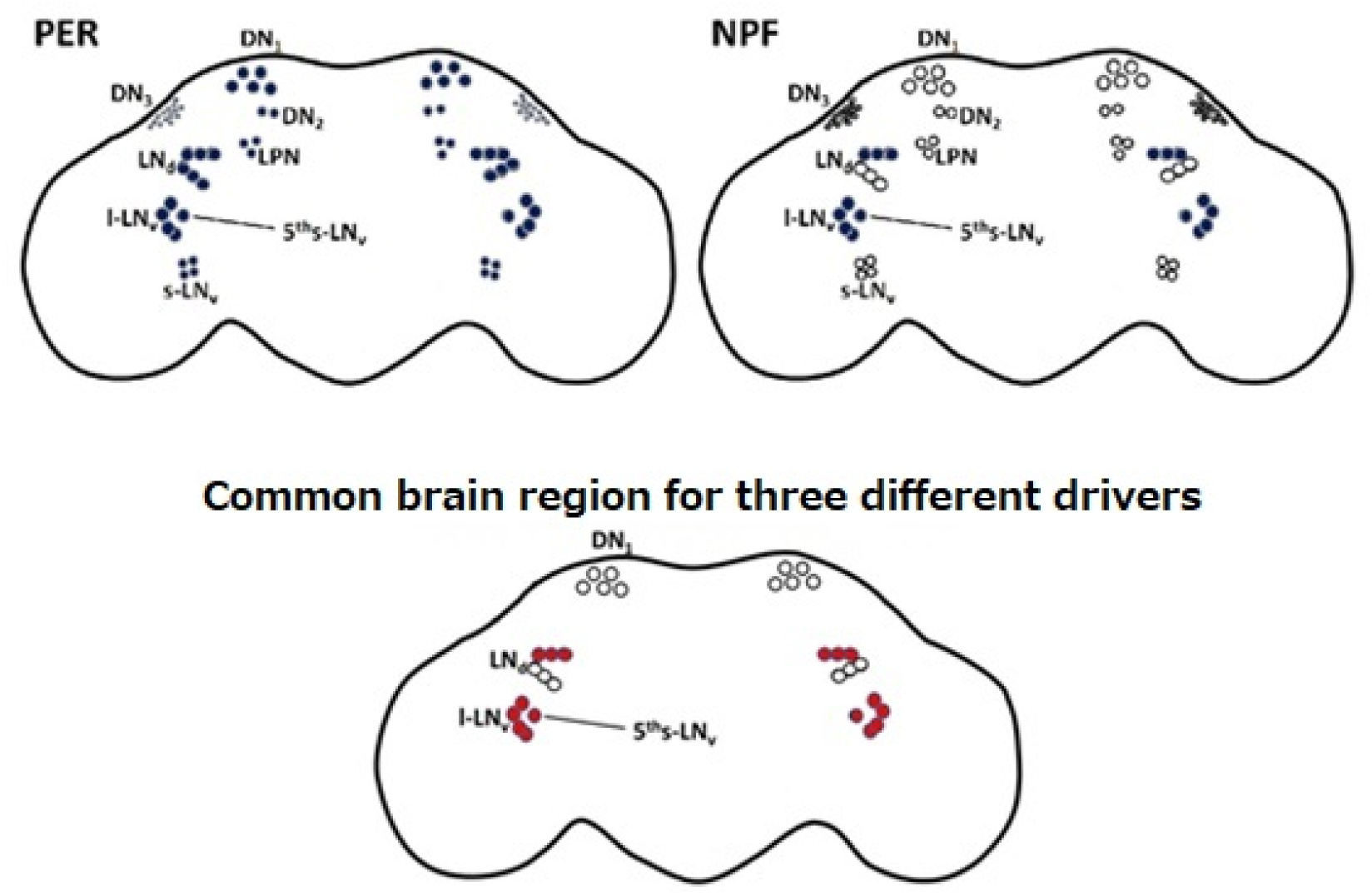
Common brain region that is necessary for the *Inos* gene expression in *Drosophila* CP rhythm. PER; Indicated cells (blue) are the expression region of *period* gene. NPF; Indicated cells (blue) are the expression region of *npf* gene. *arm* driver is expressed in all neuronal cells. Common brain region; Indicated cells (red) are the common expression region within three different promoters (*arm, per, npf*). The common brain region were shown as red (LNd, l-LNv and 5^th^ s-LNv).

## 3. Discussion

Here, we showed that the close-proximity (CP) rhythm of *D. melanogaster* courtship behavior was damped under low-nutrient conditions but enhanced by feeding the flies with *Myo*-inositol. Dietary 5% and 10% *Myo*-inositol lengthened the period of the CP rhythm. *Myo*-inositol is known to improve clinical pregnancy rates in infertile men and women by improving sperm quality and ovulation induction *in vivo* and *in vitro* (Korosi et al., 2017). Furthermore, fission yeast requires *Myo*-inositol for mating and sporulation (Niederberger et al., 1998; Voicu et al., 2002). In *D. melanogaster*, disruption of *Inos* gene causes male sterility (Jackson, Flores, Eldon, & Klig, 2018). Furthermore, *Inos* is highly expressed in testes and the head in *D. melanogaster* (Chen et al., 2014). These findings with consideration with our data suggest that *Myo*-inositol may be an important factor for the mating activity in several different species.

To determine the internal synthesis effect of *Myo*-inositol on CP rhythms, we established *Inos* gene knock down RNAi strains using the *armadillo*-GAL4, *period-*Gal4 and *neuropeptide F*-Gal4 drivers. All three *Inos* knock down strains showed arrhythmic CP behavior, but still intact for the locomotor rhythms. We also found the normal *per* gene rhythmic expression in the brains and bodies of the three RNAi lines. These results suggest that *Inos* is involved in the circadian mating rhythm in *D. melanogaster* (Fig. 5).

**Fig. 5.**
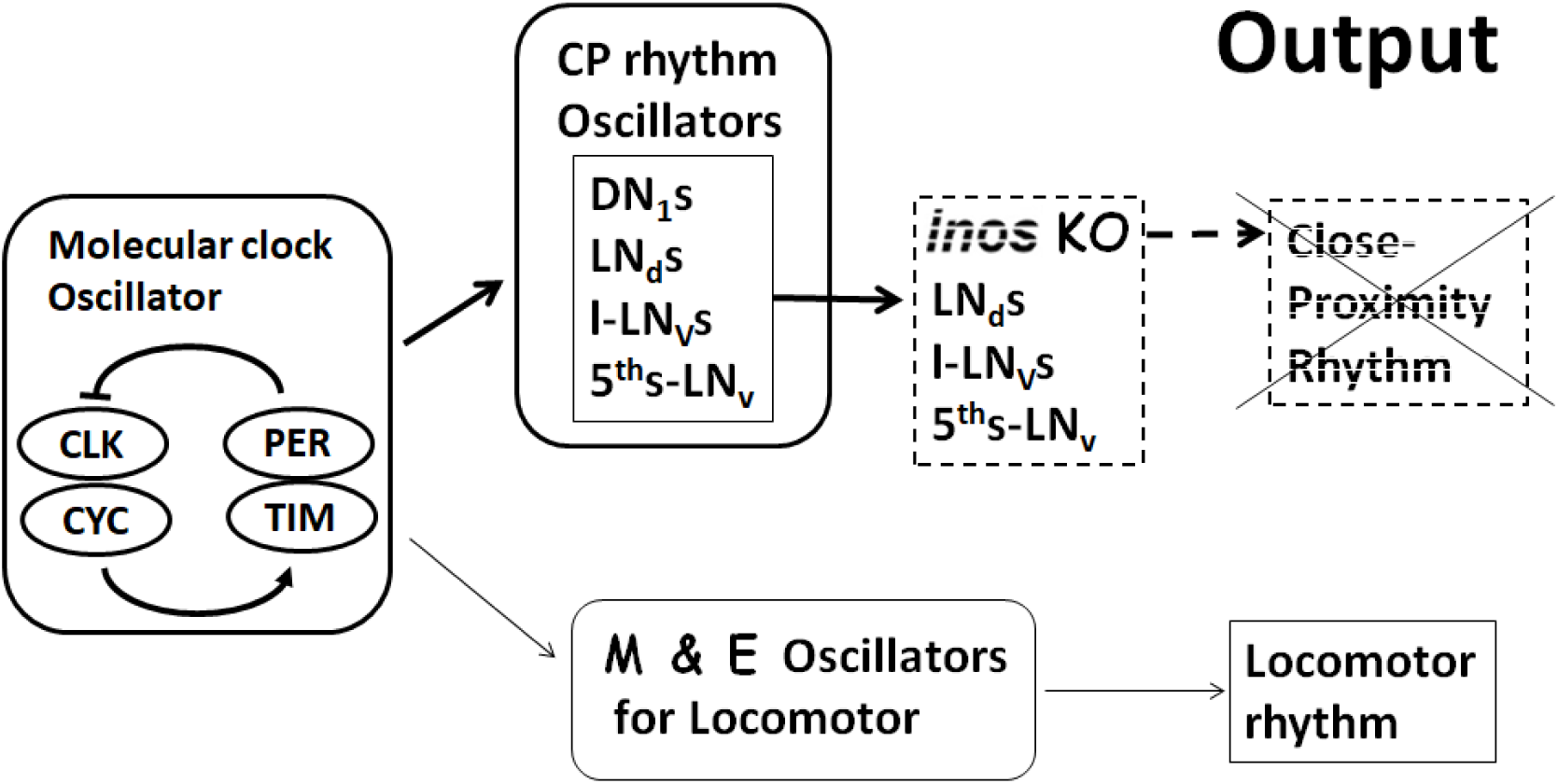
Model for CP rhythm generation. *Drosophila* with three different *Inos* RNAi lines did not alter locomotor rhythm but abolished CP rhythm. This suggests that the *Inos* gene expression in upper LN_d_ l-LN_V_ 5^th^s-LN_v_ in brain might be an output for CP rhythm oscillators and Molecular clock Oscillator. M; morning oscillator, E; evening oscillator. KO; knock out.

The summary of the brain region for three-different *Inos* knock down expression suggests that *Inos* gene expressions of upper LN_d_, l-LN_V_ 5^th^s-LN_v_ in brain are necessary for the proper CP rhythm generation in *D. melanogaster* (Fig. 3, 4). To clarify such conclusion, the region-specific *Inos* gene expression under *Inos* null back ground is required.

Previously, the nuclei for circadian CP rhythm generation (CP rhythm oscillators) using the disruption of molecular clock oscillator cells have been reported differently with locomotor oscillator by us and others (Fig. 4) (Fujii, Emery, & Amrein, 2017; Fujii et al., 2007; Hamasaka et al., 2010). Comparing with these data, DN1s are not required for CP rhythm generation in *Inos* gene expression (Fig. 4, 5). Thus, we hypothesized the *Inos* gene expression in upper LN_d_, I-LN_V_ 5^th^s-LN_v_ of fly brain might be important as an output for CP rhythm oscillators in this paper (Fig. 4,5) (Fujii et al., 2007; Hamasaka et al., 2010; Sakai & Ishida, 2001). In fact, we found circadian *Inos* mRNA expression (data not shown) in whole body and several E-box sequences on *Inos* geneomic sequence in *D. melanogaster*. These findings also suggest that *Inos* is out of output genes for the molecular circadian clock oscillator loop including CLK, CYC, PER, and TIM (Fig. 5).

In this paper, dietary *Myo*-inositol and the knock down of internal *Myo*-inositol synthesis showed different effects to the circadian mating behavior and the locomotor behavior. We think the tissue of action for dietary *Myo*-inositol might be different with that of CP rhythm generation in brain. A more precise molecular biological study is required to support these notions.

## 4. Experimental procedures

### 4.1. Fly food preparation

Fly food was prepared as described previously (Sakata et al., 2015). Boiled standard medium consisting of 8% corn meal, 5% glucose, 5% dry yeast extract, 0.64% agar was supplemented with 0.5% propionic acid and 0.5% butyl p-hydroxybenzoate (Standard food, SF). Designated low-nutrient food (LNF) comprising 5% glucose, 1.5% agar, 0.5% butyl *p-*hydroxybenzoate was supplemented without or with *Myo*-inositol (1%, 4% or 10%). *Myo*-inositol was commercially available (Wako Pure Chemical Industries Ltd, Osaka, Japan).

### 4.2. Close-proximity assays

More than 40 male and female flies were maintained in different vials with SF for three days starting from the third day after eclosion. One male and one female from the same genotype were lightly anesthetized with CO_2_ and rapidly placed in 35-mm-diameter dishes containing SF or LNF. The dishes were then mounted under a CCD camera, (Watec Co. Ltd., Yamagata, Japan) which is sensitive to light at the near infra-red range and a recording system was established as described (Fujii et al., 2007; Hamasaka et al., 2010). A fluorescent lamp provided illumination at 100 lux and a red LED provided constant dim light < 1 lux. Time-lapse images (1 frame per 10 s) were sent to a personal computer. The locations of the flies on the X and Y-axes of the images were determined using ImageJ Plugin (http://rsb.info.nih.gov/ij/). The CP index of each pair was calculated from the X-Y value with a threshold (< 5 mm) between them. Male flies moving to within 5 mm of a female and those remaining > 5 mm from a female were scored as 1 or 0, respectively, in the algorithm of the CP index program. All CP assays proceeded with flies of the same genotypes and the data were averaged for each genotype. The circadian rhythmicity of CP was determined using autocorrelation (CORREL function) analysis (Sakata et al., 2015). The free-running period and the power of rhythmicity in each genotype were calculated as the average of the free-running period and the maximum correlation between each pair evaluated by autocorrelation as being rhythmic (CORREL function) (Hamasaka et al., 2010). These all steps were automatically recorded by AUTOCIRCAS (Taisei Co. Ltd. Chiba, Japan).

### 4.3 Locomotor assays

Locomotor assays were performed as described previously (Nishinokubi, Shimoda, & Ishida, 2006). We tracked the movements of flies that were individually housed with medium, using infrared sensors and a *D. melanogaster* activity monitor (Trikinetics Inc, Waltham, MA) placed in an incubator under DD at 25°C ± 0.5°C. Male flies were assessed at 3 days after eclosion. Signals from the sensors were summed every minute using a computer. Flies were entrained by daily cycles of 12□h light followed by 12□h complete darkness (LD12). After 2 days of entrainment, flies were exposed to constant dark (DD).

### 4.4. Fly rearing and crosses

*D. melanogaster* strains were maintained as described previously (Sakata et al., 2015). Flies were reared in vials of standard yeast cornmeal at 25°C and entrained to 12 h light and dark periods (LD 12:12h). The wild-type strain (Oregon R), clock mutant (*per*^*0*^) and transgenic strain (*UAS-Inos*^*RNAi*^, *armadillo-Gal4, period-Gal4* and *neuropeptide F-Gal4*) were used for evaluating CP rhythm in this experiment. Schema of crosses to obtain transgenic flies is shown in Fig. S2. *D. melanogaster* strains were obtained from Bloomington *Drosophila* Stock Center (http://fly.bio.indiana.edu/) except for *Inos* RNAi lines (Vienna Drosophila Resource Center; https://stockcenter.vdrc.at/control/main).

### 4.5. Isolation of RNA and quantitative RT-PCR

Flies were entrained at 25 °C under 12 h light and dark periods (LD 12:12h) and then male, five-day-old whole nine flies were homogenized in TRIzol reagent (Invitrogen, Carlsbad, CA, USA), mixed with 25% chloroform and separated by centrifugation at 12,000 ×g for 15 min in 4 °C.

Supernatants were mixed with an equal volume of 2-propanol, separated by centrifugation at 12,000g for 10 min at 4 °C and then the pellets were mixed with 70% ethanol, separated by centrifugation at 7500g for 5 min at 4 °C and mixed with dH_2_O. Complementary DNAs were synthesized using the Prime Script RT Reagent Kit (Takara Bio, Otsu, Japan) according to the manufacturer’s protocol.

The amounts of cDNA generated from *Inos* and *RpL32* genes were measured by quantitative RT-PCR using a Light Cycler Nano (Roche Applied Science) and FastStart Essential DNA Green Master (Roche Applied Science). The amount of mRNA was corrected relative to that of *RpL32*.

Primer sequences for *Inos* were *Inos-F*: TTTTCATTGCCGGCGATG; *Inos-R*: TGTAACTGGCAATGGACACCG and those of *RpL32*: *RpL32-F*: AGATCGTGAAGAAGCGCACCAAG; *RpL32-R*: CACCAGGAACTTCTTGAATCCGG.

### 4.6. Statistical analysis

All CP data were averaged for each genotype. The circadian rhythmicity of the CP was determined using autocorrelation analysis (Hamasaka et al., 2010; Levine, Funes, Dowse, & Hall, 2002). The free-running period and the power of rhythmicity in each genotype were calculated as the average of the free-running period. The maximum correlation value of each pair was evaluated for the rhythmicity from the autocorrelation analysis. The period and number of rhythmic pairs were statically analyzed using a two-tailed t-test. Data are expressed as means ± SEM.

## Contributions

N.I. designed the research; K.S. and H.K. performed research; H.K. analyzed data; and N.I. and H.K. wrote the paper.

## Supplemental information

**Fig. S1.**
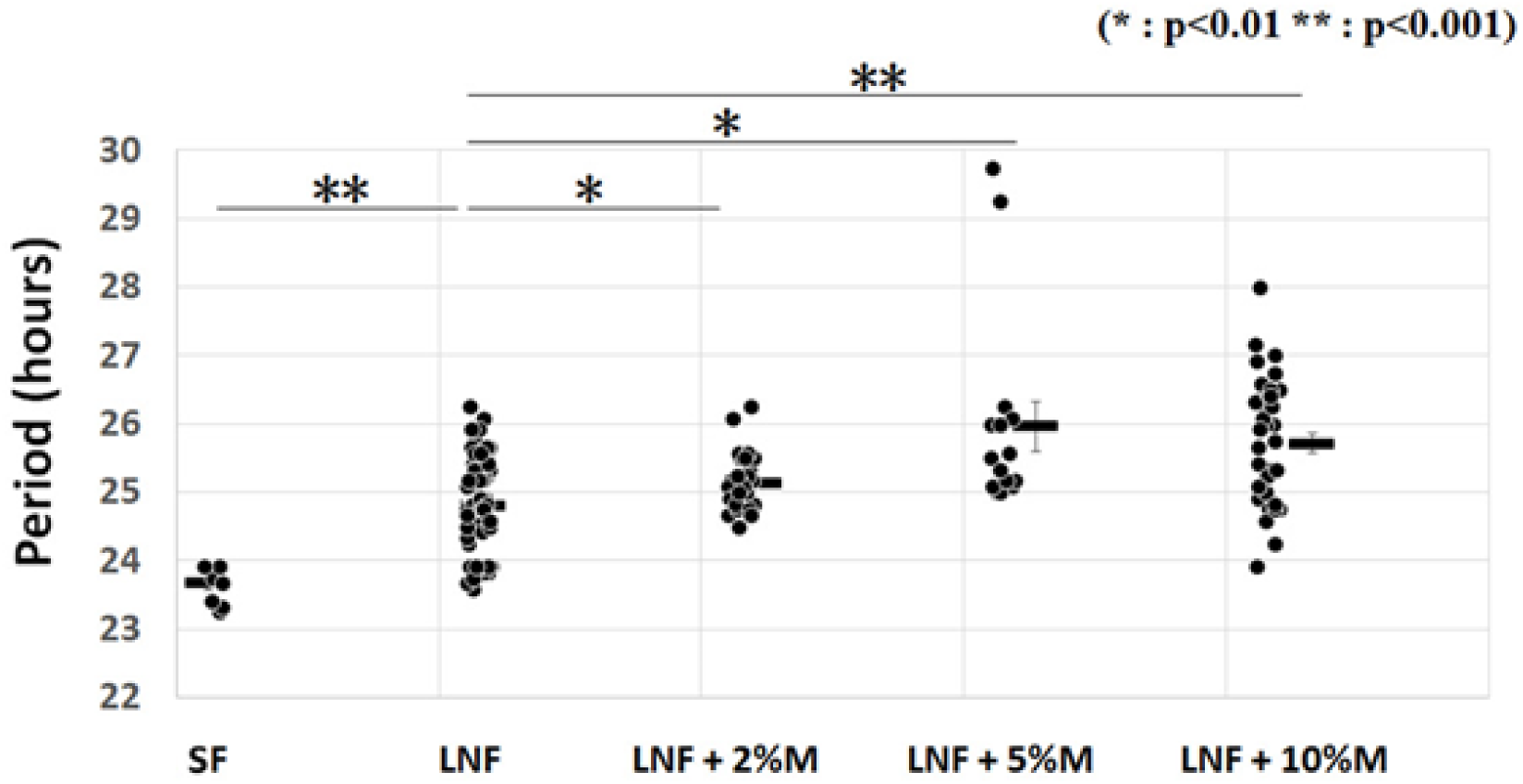
Dietary *Myo*-inositol lengthens the wild-type *Drosophila* locomotor rhythms. The period of wild-type (Oregon R) fly locomotor rhythms were increased by dietary *Myo*-inositol in LNF under low-nutrient food. Data are shown as means ± SE. *Significant difference compared with control (*p < 0.01; **p < 0.001; t-test). SF: Standard food, LNF: Low-nutrient food.

**Fig. S2.**
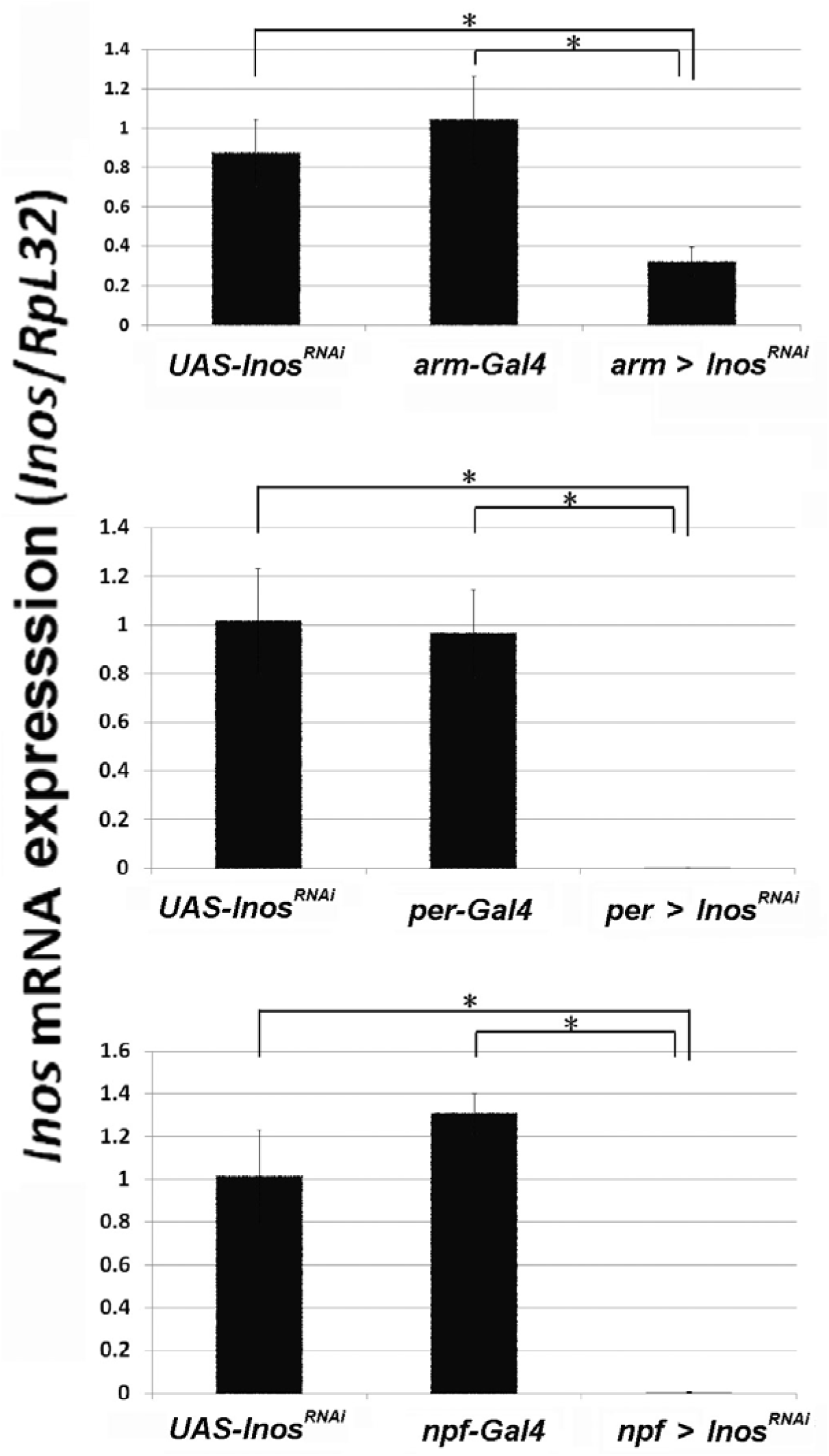
Three different RNAi strains decreased *Inos* gene expression. *Inos* RNA expression of RNAi flies were measured by qRT-PCR. *Inos* gene expression was significantly reduced by using *armadillo, period*, and *neuropeptide F* –Gal4 drivers. Data are shown as means ± SE. *Significant difference comparedwith control (*p < 0.05; t-test).

**Fig. S3.**
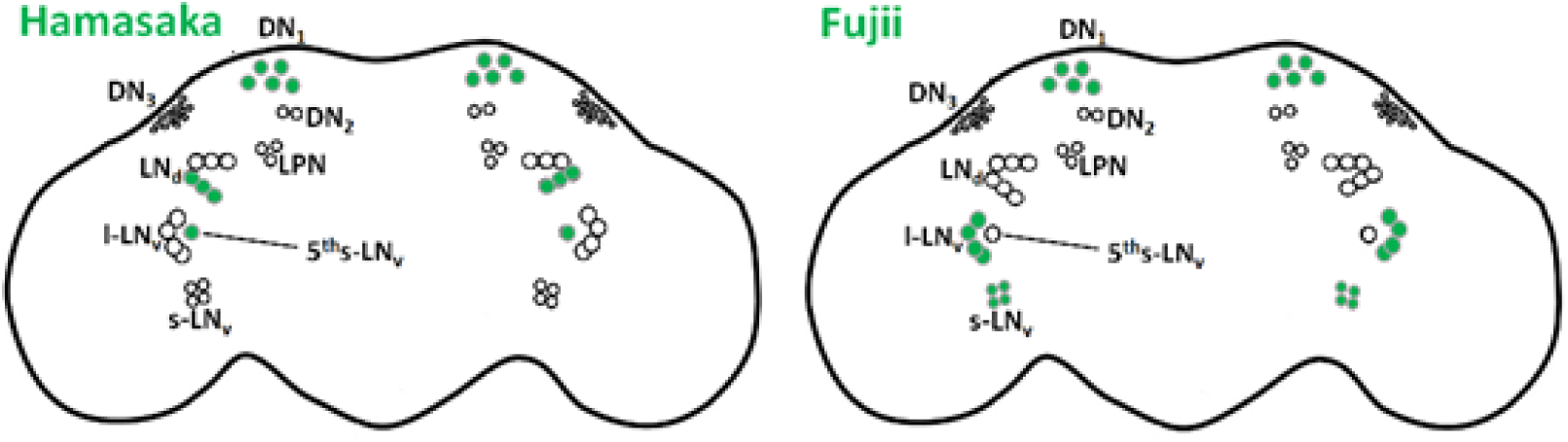
Shematic diagram of CP rhythm oscillator cells in *Drosophila* brain. Green indicates CP rhythm osillator cells necessary for CP rhythm generation reported in our previous study (left; Hamasaka et al., 2010) and reported by Amrein’s group (right; Fujii et al., 2017).

